# An Evaluation of Machine-learning for Predicting Phenotype: Studies in Yeast, Rice, and Wheat

**DOI:** 10.1101/105528

**Authors:** Nastasiya F. Grinberg, Oghenejokpeme I. Orhobor, Ross D. King

## Abstract

In phenotype prediction, the physical characteristics of an organism are predicted from knowledge of its genotype and environment. Such studies are of the highest societal importance, as they are of central importance to medicine, crop-breeding, etc. We investigated three phenotype prediction problems: one simple and clean (yeast), and the other two complex and real-world (rice and wheat). We compared standard machine learning methods (elastic net, ridge regression, lasso regression, random forest, gradient boosting machines (GBM), and support vector machines (SVM)), with two state-of-the-art classical statistical genetics methods (including genomic BLUP). Additionally, using the clean yeast data, we investigated how performance varied with the complexity of the biological mechanism, the amount of observational noise, the number of examples, the amount of missing data, and the use of different data representations. We found that for almost all phenotypes considered standard machine learning methods outperformed the methods from classical statistical genetics. On the yeast problem, the most successful method was GBM, followed by lasso regression, and the two statistical genetics methods; with greater mechanistic complexity GBM was best, while in simpler cases lasso was superior. When applied to the wheat and rice studies the best two methods were SVM and BLUP. The most robust method in the presence of noise, missing data, etc. was random forests. The classical statistical genetics method of genomic BLUP was found to perform well on problems where there was population structure, which suggests one way to improve standard machine learning methods when population structure is present. We conclude that the application of machine learning methods to phenotype prediction problems holds great promise.

## 1 Introduction and Background

### 1.1 Predicting Phenotype

The phenotype (physical character) of an organism is the result of interactions between the organism’s complement of genes (its genotype) (Wei et al., 2014; Mackay, 2014) and its environment. A central problem of genetics is to predict an organism’s phenotype from knowledge of its genotype and environment. This problem is now of the highest societal importance. For example, human disease is a phenotype and understanding its relation to genotype and environment is a central problem in medicine (Stranger et al., 2011; Lee et al., 2011), see for example the recent studies in schizophrenia (Schizophrenia Working Group of the Psychiatric Genomics Consortium, 2014), obesity (Locke et al., 2015), educational achievement (Rietveld et al., 2013), etc. Similarly, crop yield and drought resistance are phenotypes, and if we are going to be able to feed the world’s growing population, it is essential to better predict crop phenotypes from knowledge of their genotypes and environment (Buckler et al., 2009; Jannink et al., 2010; Hayes and Goddard, 2010; Brachi et al., 2011; Desta and Ortiz, 2014). For such reasons the problem of predicting phenotype has recently (2016) been listed by the US National Science Foundation as one of its six key ‘Research Frontiers’: ‘understanding the rules of life’.

Our ability to predict phenotype is being revolutionised by advances in DNA sequencing technology. These advances have enabled, for the first time, an organism’s genotype to be extensively characterised, typically via thousands of genetic markers. The cost of sequencing is decreasing rapidly, which means that it is now often low enough that in a single investigation many organisms (hundreds/thousands) may be genotyped, which opens up the possibility of using statistics/machine learning to learn predictive relationships between an organism’s genotype, environment, and phenotype. Such studies are often called Genome-Wide Association Studies (GWAS).

The traditional focus of most GWAS has been on the discovery of genetic markers (normally only a small number) that are ‘associated’ (i.e., correlated) with a phenotype. This has limitations. The focus on a small number of genes has significant biological limitations as most biological phenotypes result from the interaction of multiple genes and the environment. The focus on association rather than prediction has statistical limitations as it makes the objective evaluation of the utility of results difficult.

The current trend is therefore towards a more direct and operational approach to phenotype prediction problems: learn a predictive function that, from the input of an organism’s genotype and environment, predicts its phenotype (see, for example, Yang et al., 2010; Bloom et al., 2013; Desta and Ortiz, 2014; Shigemizu et al., 2014). Special purpose statistical genetics methods have been developed for this task (see, for example, Lynch and Walsh, 1998; Bloom et al., 2013; Desta and Ortiz, 2014). Predictive phenotype problems are also clearly well suited for standard machine learning methods.

In this paper, we compare for phenotype prediction a state-of-the-art classical statistical genetics method and a mixed-model approach BLUP (used extensively in genomic selection applications) with standard machine learning methods. We investigate how these methods perform on three very different types of phenotype prediction problem, one from yeast *Saccharomyces cerevisiae* (Bloom et al., 2013), the other two from wheat *Triticum aestivum L.* (Poland et al., 2012) and rice (Alexandrov et al., 2015). We also compare how performance varies with the complexity of the biological mechanism, the amount of observational noise, the number of examples, the amount of missing data, and the use of different data representations.

### 1.2 Phenotype Prediction Data

Genomic data has a specific structure that strongly influences the application of machine learning methods. We therefore first briefly describe this structure using standard machine learning terminology—rather than the terminology of statistical genetics which may confuse the uninitiated.

#### Representing genotype

The complete genotype of an organism consists of the linear arrangement of its genes on chromosomes, together with the sequences of all the genes and intergenic regions. Genotype information is normally represented in phenotype prediction problems as ‘markers’, these are discrete attributes that signify that a particular stretch of DNA varies between organisms. Usually, these variations are mutations at a single position (base pair) in a DNA sequence called single-nucleotide polymorphisms (SNPs). As organisms of the same species mostly share the same DNA sequences, this representation is reasonably concise, but as genomes are large, many thousands of markers are typically needed to characterise an organism’s genome. (It should be noted that this propositional marker representation is sub-optimal, as it ignores a substantial amount of information: the biological context of the markers involved, the linear ordering of the markers, etc.) In this paper we utilise this marker representation as it is standard, and it simplifies the application of standard statistical/machine learning methods.

In phenotype prediction problems it is preferable for all the organisms (examples) to have their genotypes fully sequenced, as this provides the maximum amount of information. However, this is not possible in many problems, either because of technical reasons or cost. In such cases, the genotypes are not fully characterised. In this paper, we investigate two problems (yeast and rice) where all the organisms are fully sequenced, and another (wheat) where only a subset of markers is known, and the organism’s genome has not yet been sequenced because of its complexity.

#### Environment

Prediction of phenotype is easier if the environment is controlled. However, this is difficult or impossible to do in many cases, for example, in studies involving humans many aspects of the environment are unknown, in outdoor crop studies the weather cannot be controlled, etc. In this paper, we investigate one problem (yeast) where the environment is fully controlled (well-defined laboratory conditions), and another two (wheat and rice) where the environment is partially controlled.

#### Measuring phenotype

Due to the continuing steep fall in DNA sequencing costs in many phenotype prediction problems, the most expensive step is the observation of an organism’s phenotype. This means that in many scenarios the number of attributes is greater than the number of examples. Furthermore, it has also led to new genetic methodologies based on phenotype prediction, an example of which is genomic selection (Meuwissen et al., 2001; Heffner et al., 2009). In this paper we investigate one problem (yeast) where the observation of phenotype (growth) is cheap as it involves laboratory automation equipment, and another (wheat and rice) where it is expensive and time consuming—it takes many months for a harvest.

#### Causation of phenotype

The number of genetic mutations involved in causing a phenotype can vary greatly between phenotypes. For example, the phenotype of pea colour (yellow or green), that was classically studied by Gregor Mendel, is caused by variations (polymorphisms) in a single gene (stay-green) (Armstead et al., 2007). Therefore, given knowledge of markers in stay-green, one can usually very accurately predict pea colour. In contrast, the phenotype of human height (classically studied by Sir Francis Galton in the original ‘regression’ problem) involves a large number of genes and environmental effects—the central-limit theorem thereby explains why human height is roughly normally distributed when controlled for sex and geographical population (Wood et al., 2014).

An important feature of genetic data is that the examples (organisms) are formed by meiosis (sex), where parents shuffle their genomes when forming offspring. The mechanisms involved in meiosis are complicated, but the result is that each of the child cell’s chromosomes consists of a random patchwork of linear sections taken from the two parents (Figure 1a). This means that if two markers are close together on an organism’s DNA, then the associated mutations (allele types) are likely to be inherited together. Consequently, attributes (markers) can be highly correlated (linkage disequilibrium). It also means that all the markers in each of these linear sections are identical to that of one parent so that the variety of children that are likely to be formed is constrained.

**Fig. 1:**
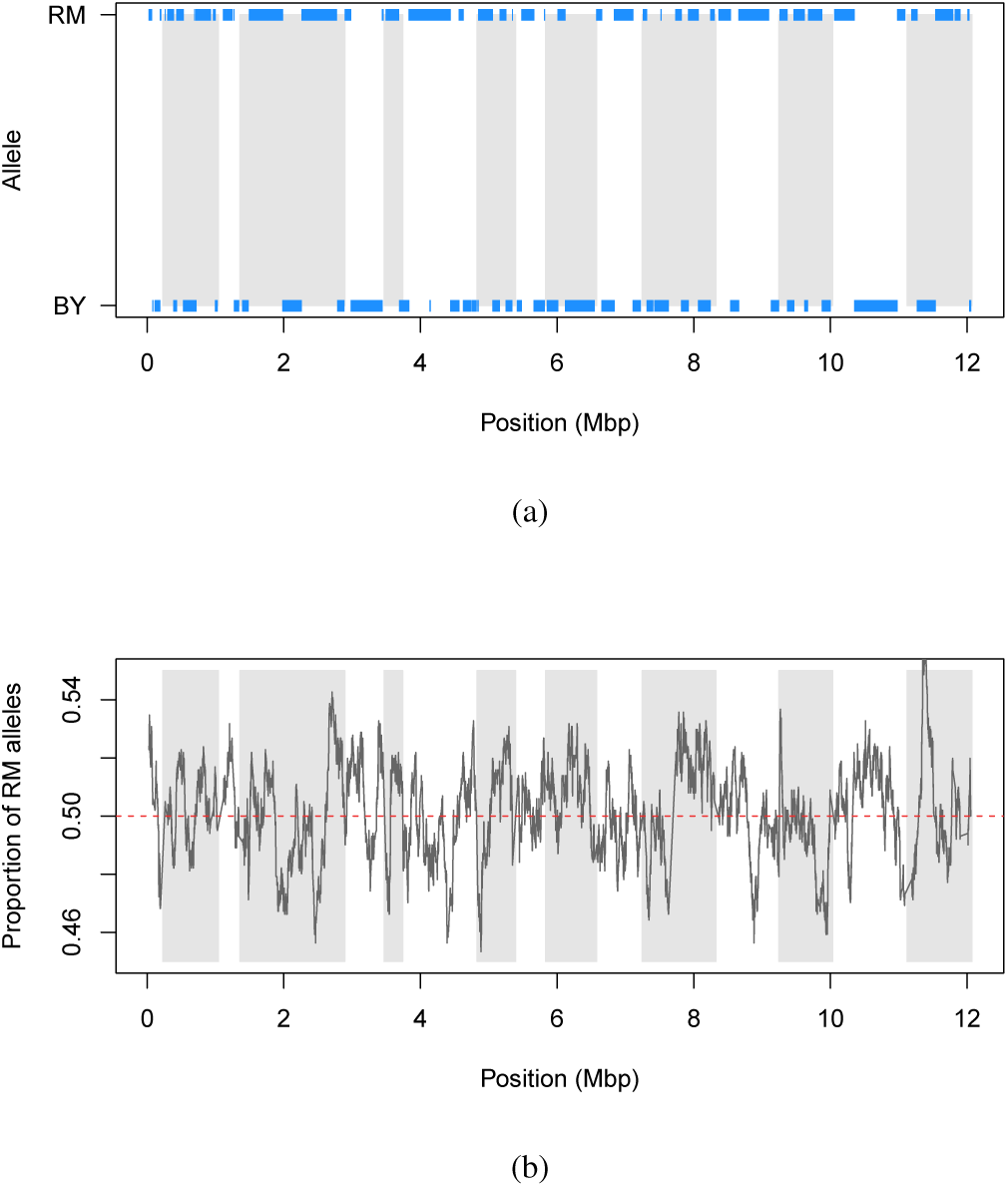
(a) Markers’ expression for a RM × BY yeast strain (segregant A01-01 (Bloom et al., 2013)) plotted against their position (mega base pairs) on the genome. Each tick represents a marker, bands represent chromosomes. 1/0 marker values correspond to RM/BY parent, respectively. (b) Proportion of markers coming from the RM parent plotted against markers position (mega base pairs) on the genome. White and grey bands separate the 16 chromosomes.

Due to meiosis populations of organisms typically have complex ancestral (pedigree) interrelationships. These interrelationships take the form of a directed acyclic graph (DAG), with meiosis events being the nodes. However, in many phenotype prediction problems the structure of the DAG is unknown. (Note that the DAG imposes a natural metric between examples (organisms) based on genetic changes resulting from meiosis.) Much of classical statistical genetics research has focused on dealing with population structure. For example, the BLUP (best linear unbiased prediction) method encodes genetic similarities between pairs of individuals in a genomic similarity matrix (GSM) (Meuwissen et al., 2001; Speed and Balding, 2014).

Many organisms have multiple genomes in each cell. Humans have single copies (haploid) in their sex cells (sperm/eggs), but otherwise have two copies (diploid), one from their father and the other from their mother. This complicates phenotype prediction as it is not clear how the genomes interact to cause phenotype. This complication is related to multi-instance learning problems (Ray and Page, 2001). In this paper we investigate one problem (yeast) where the observed organisms are haploid, i.e., there is no complication with multiple genomes. Rice is diploid, with two paired sets of genomes, while wheat is hexaploid with three pairs of paired genomes.

### 1.3 Types of Phenotype Prediction Problem

The simplest form of phenotype prediction problem is the case when a pair of parent organisms breed to produce a large set of offspring. In such cases the offspring can reasonably be assumed to be randomly generated from a given distribution: indeed the analogy with randomly dealing hands of cards is close and is commonly used in genetics (Hogben, 1946). This type of phenotype prediction problem is closely connected to practical phenotype prediction problems, for example: which embryo to select?

To investigate this type of problem we utilised a large phenotype prediction study in yeast (Bloom et al., 2013). In this study there are a large number of examples, the complete genomes of the organisms are known, the organisms are haploid, a large number of different environments were investigated under controlled laboratory conditions, and the phenotype of growth was accurately measured. Taken together these features give a phenotype predictions dataset that is as clean and complete as it is currently possible to get. Moreover, the uniform laboratory conditions under which the yeast was grown ensured that there were no (or nearly no) confounding environmental factors.

We chose as a second, and comparative, phenotype prediction problem—the real-world problem of predicting phenotype in crops: wheat (Poland et al., 2012) and rice (Alexandrov et al., 2015). This type of phenotype prediction problem is typical of genomic selection problems in organisms, in which genome-wide molecular markers are used to predict the breeding utility of individuals or breeding lines prior to phenotyping. The wheat dataset we investigated comes from a study involving 254 varieties (breeding lines) of wheat (Poland et al., 2012). These varieties were derived through six generations of meiosis (crosses) from a set of ancestor varieties. Experimental design methods were used to control the environment, and different irrigation techniques investigated. This dataset is more complex and difficult to predict than the yeast one for a number of reasons: the complete genotypes of the organisms are not known (only the markers, indeed wheat has still not been fully sequenced), the organisms are hexaploid, the organisms come from different parents (although there is some knowledge of the relationships between parents), there are fewer examples, and the environment is not fully controlled.

The rice dataset comes from the 3000 rice genomes project (Alexandrov et al., 2015). Like the wheat dataset, this problem also involves the selection of individuals that will serve as parents for the next generation of progeny using genomic predictions. The phenotype data is from different years of screening without replication. However, the values do not show significant variation due to environmental differences, as the data is part of a routine characterization of genetic resources performed by the International Rice Genebank at the International Rice Research Institute.

### 1.4 Classical Statistical-Genetics Methods for Predicting Phenotype

The most common classical statistical genetics approach to the analysis of genotype/environment/phenotype data has been to use univariate and bivariate statistical methods (Lynch and Walsh, 1998; Westfall et al., 2002; Marchini et al., 2005). These typically test each marker (or pairs of markers) for association with a phenotype individually, and independently of the other markers. The focus on such approaches seems to have been because of a desire to understand and possibly control the mechanism that causes phenotype, through identification of the markers involved. This is reasonable, but the assumption of independence does not reflect the complex causal relationships involved in phenotype formation (e.g., it ignores systems biology), and it is prone to missing markers with small effects.

The emphasis on univariate and bivariate correlations also raises the problem of multiple testing and is hindered by typical *p*-value limitations, such as dependence on sample size, minor allele frequency, and difficulty to determine a meaningful threshold for the study. The multiple testing problem may be classically addressed by using false discovery rate (FDR) (Benjamini and Hochberg, 1995) instead of the conventional *p*-values or the Bonferroni correction. One of the more recent approaches is to use Bayes factors instead of *p*-values (Wakefield, 2007), thus taking prior belief of association into account. The problem of the interdependence of hypotheses in multiple testing (that is, possible interactions between markers) has been addressed for example by using hidden Markov models (Sun and Tony Cai, 2009) and graphical models (Liu et al., 2012). In statistical genetics, arguments based on multiple testing are often used to claim that it is not possible to identify complicated interactions between markers in datasets that are not very large (Gauderman, 2002; Wang and Zhao, 2003). These arguments are incorrect as they would imply that multivariate learning is generally very difficult, which is not the case.

The emphasis in classical statistical genetics on univariate and bivariate methods research has also led to efforts to reduce the dimensionality of GWAS problems. This, for example, can be done by grouping markers in haplotypes—specific arrangements of alleles from a parent (Clark, 2004). This enables the simultaneous testing of associations between a phenotype and several markers in a target region. However identifying meaningful haplotypes is in itself a non-trivial task (Meng et al., 2003; Lin and Altman, 2004).

A change of focus from association to prediction cuts through most of the problems of the significance of associations: the validity of a multivariate relationship between markers is demonstrated by success of predictions on test data. The utility of using test data is appreciated in statistical genetics (Lynch and Walsh, 1998; Bloom et al., 2013; Desta and Ortiz, 2014; Shigemizu et al., 2014) but should perhaps be stressed more.

Recently the importance of considering all the markers simultaneously has been widely recognised (e.g. de los Campos et al. (2013)) with various multivariate linear models gaining popularity. Amongst them are genomic BLUP (Meuwissen et al., 2001; VanRaden, 2008) (a mixed-model related to ridge regression with a pre-specified penalty parameter) and other penalised regression methods (Gianola et al., 2006; De Los Campos et al., 2009; Li and Sillanpää, 2012) and Bayesian techniques (Meuwissen et al., 2001; De Los Campos et al., 2009; Habier et al., 2011; Guan and Stephens, 2011; Zhou et al., 2013). An important extension of BLUP proposed by Speed and Balding (2014) relaxes the assumption of constant variance for SNP effects. Many improved and efficient linear mixed model (LMM) algorithms have also been introduced in recent years (Kang et al., 2008; Zhang et al., 2010; Lippert et al., 2011; Loh et al., 2015; Lee and van der Werf, 2016), some of which are capable of dealing with several correlated phenotypic traits (Korte et al., 2012; Zhou and Stephens, 2014; Casale et al., 2015) (see also Widmer et al. (2014) and references therein). The attractiveness of these linear techniques lies in the fact that they take population structure and genetic relatedness into account (Price et al., 2010). However, most of these techniques have difficulty accounting for interactions.

### 1.5 The Applicability of Machine Learning Methods

Standard off-the-shelf machine learning methods (Dudoit et al., 2002; Ziegler et al., 2007; Szymczak et al., 2009; Ogutu et al., 2011, 2012; Pirooznia et al., 2012; Mittag et al., 2012; Okser et al., 2014; Leung et al., 2016) present an attractive alternative to classical statistical genetics methods. These machine learning methods are easy to use, are freely available in a variety of implementations, and intrinsically multivariate. They also do not require assumptions about the genetic mechanism underlying a trait in question (e.g., additivity of effects, the number and size of interactions, depth of interactions, etc.). In addition, machine learning methods are available that perform attribute selection (e.g., lasso and regression trees). There are also machine learning methods available that can identify complex interactions between attributes (e.g., random forest, boosted trees, neural nets), not simply bivariate ones. Furthermore, in typical GWAS problems there is the *p* ≫ *n* problem (where the number of attributes (*p*) greatly exceeds the number of sample points (*n*)), and this is a problem for classical multivariate regression, but less so for many machine learning methods.

## 2 Materials and Methods

### 2.1 Experimental Data

The yeast dataset was derived from a study of 1,008 haploid yeast strains derived from a cross (meiosis) between a laboratory and a wine strain of the yeast *Saccharomyces cerevisiae*. The parent strains differed by 0.5% at the sequence level. The genotypes of the parents and children were determined by sequencing (Figure 1). The raw sequence data was transformed into 11,623 Boolean markers: coded to be ‘1’ if the sequence variation came from the wine strain (RM) parent and ‘0’ if it came from the laboratory strain (BY) parent.

The environment of the yeast strains was modified in 46 different ways (Table 1): the basic chemicals used for growth were varied (e.g., galactose, maltose), minerals added (e.g., copper, magnesium chloride), herbicides added (paraquat), etc.

**Table 1:** cv*R*^2^ for the five ML methods, BLUP and for the QTL mining approach of Bloom *et al.* The best performance for each trait is in boldface.

| Trait/Method | Bloom et al | Lasso | Ridge | BLUP | GBM | RF | SVM |
| --- | --- | --- | --- | --- | --- | --- | --- |
| Cadmium Chloride | 0.780 | 0.779 | 0.445 | 0.556 | **0.797** | 0.786 | 0.565 |
| Caffeine | 0.197 | 0.203 | 0.171 | 0.229 | **0.250** | 0.236 | 0.234 |
| Calcium Chloride | 0.198 | 0.255 | 0.240 | **0.268** | 0.261 | 0.205 | 0.261 |
| Cisplatin | 0.297 | 0.319 | 0.253 | 0.290 | **0.338** | 0.275 | 0.272 |
| Cobalt Chloride | 0.431 | 0.455 | 0.431 | 0.457 | **0.460** | 0.398 | 0.448 |
| Congo Red | 0.460 | **0.504** | 0.469 | 0.500 | 0.500 | 0.398 | 0.487 |
| Copper | 0.405 | 0.345 | 0.284 | 0.331 | **0.456** | 0.406 | 0.338 |
| Cycloheximide | 0.466 | 0.498 | 0.480 | 0.516 | 0.513 | 0.444 | **0.529** |
| Diamide | 0.417 | 0.479 | 0.468 | **0.498** | 0.460 | 0.318 | 0.486 |
| E6.Berbamine | 0.380 | 0.403 | 0.372 | 0.399 | **0.412** | 0.281 | 0.390 |
| Ethanol | 0.486 | 0.495 | 0.434 | 0.460 | **0.518** | 0.475 | 0.455 |
| Formamide | 0.310 | 0.238 | 0.179 | 0.232 | **0.350** | 0.298 | 0.240 |
| Galactose | 0.201 | 0.183 | 0.171 | 0.211 | **0.235** | 0.219 | 0.217 |
| Hydrogen Peroxide | 0.362 | 0.377 | 0.308 | 0.365 | 0.385 | 0.355 | **0.399** |
| Hydroquinone | 0.135 | 0.201 | 0.173 | **0.225** | 0.212 | 0.191 | 0.188 |
| Hydroxyurea | 0.232 | 0.303 | 0.266 | 0.303 | **0.336** | 0.243 | 0.320 |
| Indoleacetic Acid | **0.480** | 0.302 | 0.239 | 0.301 | 0.476 | 0.458 | 0.310 |
| Lactate | 0.523 | **0.568** | 0.522 | 0.552 | 0.561 | 0.510 | 0.557 |
| Lactose | 0.536 | 0.567 | 0.532 | 0.562 | **0.582** | 0.530 | 0.565 |
| Lithium Chloride | 0.642 | 0.704 | 0.635 | 0.670 | **0.711** | 0.538 | 0.680 |
| Magnesium Chloride | **0.278** | 0.229 | 0.196 | 0.250 | 0.266 | 0.255 | 0.267 |
| Magnesium Sulfate | **0.519** | 0.369 | 0.326 | 0.360 | 0.492 | 0.434 | 0.378 |
| Maltose | 0.780 | 0.620 | 0.488 | 0.534 | **0.809** | 0.806 | 0.522 |
| Mannose | 0.230 | 0.202 | 0.162 | 0.210 | **0.255** | 0.234 | 0.215 |
| Menadione | 0.388 | 0.412 | 0.375 | 0.407 | **0.432** | 0.396 | 0.402 |
| Neomycin | 0.556 | **0.614** | 0.580 | 0.609 | 0.600 | 0.487 | 0.597 |
| Paraquat | 0.388 | **0.496** | 0.447 | 0.474 | 0.488 | 0.298 | 0.479 |
| Raffinose | 0.317 | 0.357 | 0.341 | 0.371 | **0.383** | 0.368 | 0.364 |
| SDS | 0.348 | **0.411** | 0.345 | 0.386 | 0.393 | 0.337 | 0.383 |
| Sorbitol | **0.424** | 0.369 | 0.296 | 0.333 | 0.379 | 0.383 | 0.318 |
| Trehalose | 0.489 | 0.500 | 0.463 | 0.487 | **0.515** | 0.472 | 0.477 |
| Tunicamycin | 0.492 | 0.605 | 0.586 | 0.618 | 0.618 | 0.385 | **0.634** |
| x4-Hydroxybenzaldehyde | 0.442 | 0.411 | 0.325 | 0.365 | **0.471** | 0.404 | 0.355 |
| x4NQO | 0.604 | 0.612 | 0.487 | 0.538 | **0.636** | 0.559 | 0.542 |
| x5-Fluorocytosine | 0.354 | 0.386 | 0.321 | 0.354 | **0.397** | 0.334 | 0.373 |
| x5-Fluorouracil | 0.503 | **0.552** | 0.512 | 0.545 | 0.536 | 0.454 | 0.546 |
| x6-Azauracil | 0.258 | 0.298 | 0.270 | 0.308 | **0.315** | 0.289 | 0.279 |
| Xylose | 0.475 | 0.468 | 0.431 | 0.465 | **0.516** | 0.484 | 0.460 |
| YNB | 0.508 | 0.541 | 0.481 | 0.519 | **0.543** | 0.411 | 0.525 |
| YNB:ph3 | 0.151 | 0.18 | 0.144 | **0.195** | 0.194 | 0.153 | 0.166 |
| YNB:ph8 | 0.295 | 0.345 | 0.315 | **0.356** | 0.354 | 0.267 | 0.334 |
| YPD | 0.533 | 0.546 | 0.480 | 0.515 | **0.556** | 0.469 | 0.524 |
| YPD:15C | **0.432** | 0.383 | 0.311 | 0.345 | 0.427 | 0.424 | 0.333 |
| YPD:37C | **0.711** | 0.653 | 0.576 | 0.606 | 0.691 | 0.686 | 0.603 |
| YPD:4C | 0.406 | 0.430 | 0.396 | 0.418 | **0.485** | 0.405 | 0.421 |
| Zeocin | 0.465 | 0.469 | 0.450 | 0.482 | **0.495** | 0.360 | 0.475 |

Yeast population growth (growth is the most basic of all phenotypes) was measured under these different conditions. As the data was generated by high-throughput robotics there are many missing values; there are, for example, only 599 readings available for sorbitol. Most traits, however, have upwards of 900 readings, some with two replications (which we average). All the growth measurements are normalised to have a mean of 0 and variance of 1.0.

Using this yeast study, we investigate different aspects of applying machine learning to phenotype prediction data by starting with as clean data as possible, and then gradually artificially degrading it to make it resemble different practical phenotype prediction problems in animals and plants. A complementary motivation for using such clean and complete data is that with improved technology applied phenotype prediction problems will increasingly resemble this clean, comprehensive form.

The wheat dataset comes from a genomic selection study in wheat involving 254 breeding lines (samples) with genotypes represented by 33,516 SNP markers coded as {–1, 0, 1} to correspond to the *aa, aA* and *AA* alleles, respectively (Poland et al., 2012). Missing values were imputed with heterozygotes *aA* (the original paper found little difference between four different imputation methods, one of which was imputing with heterozygotes). The wheat lines were evaluated in plots for four phenotypic traits: yield (drought), yield (irrigated), thousand kernel weight (TKW) and days to heading (DTH). Phenotypic values were once again normalised.

For the rice data, the Core SNP subset of 3000 Rice Genomes version 0.4 from SNP-SEEK (Mansueto et al., 2016) was used. The genotypes for the samples in the dataset originally contained 996,009 SNP markers. However, a subset of 101,595 markers was used in this study to reduce computational complexity. These markers were selected by linkage disequilibrium in Plink (Purcell et al., 2007), using the --`indep-pairwise` command with a window of 50 SNPs, a step size of 5, and *r*^2^ value of 0.02. The markers were coded in the same way as in the wheat dataset, and missing values were imputed using column means. Twelve phenotypic traits were considered: culm diameter, culm length, culm number, grain length, grain width, grain weight, days to heading, ligule length, leaf length, leaf width, panicle length and seedling height. Due to missing phenotype data, each trait has its own set of samples, with the number of samples ranging from 1877 to 2265.

In statistical/machine learning terms: each of the different genotype/phenotype combinations represents a different regression problem. The yeast strains/wheat/rice samples are the examples, the markers in the examples are the attributes, and the growth of the strains (for yeast) and agronomic traits evaluated (for wheat and rice) are variables to be predicted.

## 3 Learning Methods

### 3.1 Standard Statistical and Machine Learning Methods

We investigated several variants of penalised linear regression: elastic net (Zou and Hastie, 2005), ridge regression (Hoerl and Kennard, 1970), and lasso regression (Tibshirani, 1996). The rationale for choosing these methods is that they most closely resemble the multivariate approaches used in classical statistical genetics. We also investigated an array of models that interpolate between ridge and lasso regressions through use of an elastic net penalty. We considered 11 values of the elastic net penalty *α* evenly spaced between 0 (ridge) and 1 (lasso) with the value of the overall penalty parameter *λ* chosen by cross-validation separately for each value of *α* (see Figure 2).

**Fig. 2:**
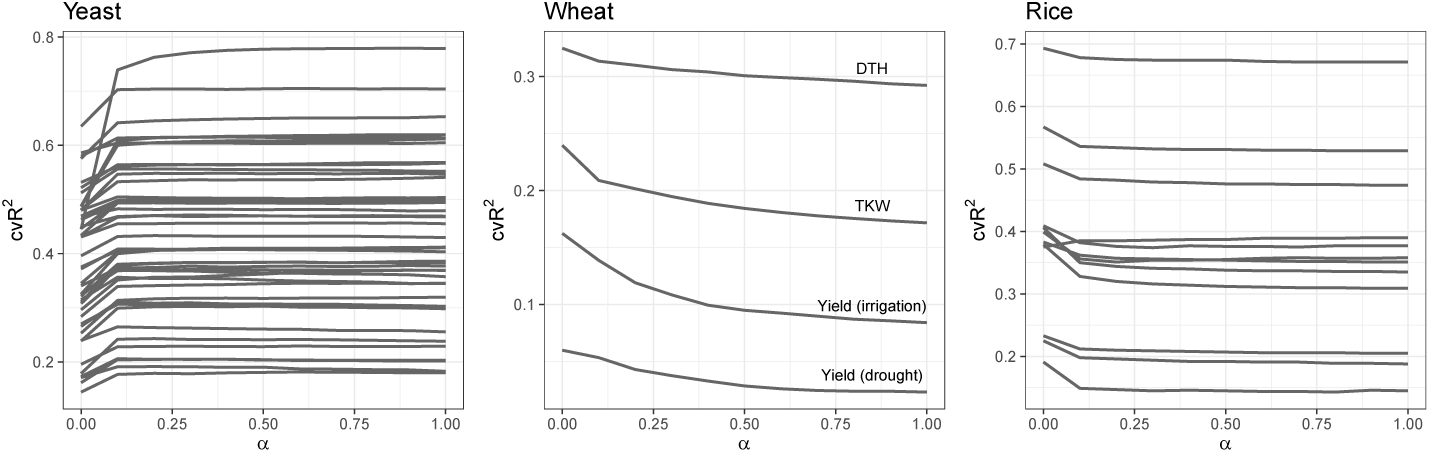
Comparison of performance of elastic net on yeast, wheat and rice datasets for varying values of the *α* parameter for each trait with *α* on the *x*-axis and cv*R*^2^ on the *y*-axis.

We also investigated the tree methods of random forests (Breiman, 2001) and gradient boosting machines (GBM) (Friedman, 2001). The rationale for the use of these is that they are known to work robustly, and have an inbuilt way of assessing the importance of attributes. For random forests we used 700 iterations for yeast and wheat and 1000 iterations for rice (chosen to be enough for convergence) of fully-grown trees with the recommended values of *p/*3 (where *p* is the number of attributes) for the number of splitting variables considered at each node, and 5 examples as the minimum node size. For GBM we tuned two parameters via internal train/test split inside each fold: interaction depth and shrinkage. We investigated interaction depths of 1, 2 and 3 and shrinkage of 0.001, 0.01 and 0.1 (in our experience the default shrinkage parameter of 0.001 lead to too slow of a convergence), resulting in a two-dimensional parameter grid. We used 1,000 trees, which was enough for convergence for all traits. The optimal number of iterations was determined via an assessment on an internal validation set within each cross-validation fold. Finally, we used the default value of 0.5 as the subsampling rate.

Finally, we investigated support vector machines (SVM) (Cortes and Vapnik, 1995). SVM methods have been gaining popularity in phenotype prediction problem recently. However, experience has shown that they need extensive tuning (which is unfortunately extremely time consuming) to perform well (Hsu et al., 2008). We used *ε*-insensitive regression with Gaussian kernel and tuned the model via internal testing within each cross-validation fold over a fine grid (on the logarithmic scale) of three parameters: *ε*, cost parameter *C* and *γ* (equal to 1/(2*σ* ^2^), where *σ* ^2^ is the Gaussian variance).

All analysis was performed in R (R Core Team, 2018) using the following packages: GLMNET for elastic net, RANDOMFOREST for random forest, GBM for gradient boosting and KERNLAB for support vector machines. The CARET package was used for tuning GBM and SVM.

### 3.2 Classical Statistical Genetics

To compare the machine learning methods with state-of-the-art classical genetics prediction methods we reimplemented the prediction method described in the original yeast Nature paper (Bloom et al., 2013), and applied the genomic BLUP model. The ‘Bloom’ method has two steps. In the first, additive attributes are identified for each trait in a four-stage iterative procedure, where at each stage only markers with LOD significance at 5% false discovery rate (identified via permutation tests) are kept and added to a linear model; residuals from this model are then used to identify more attributes in the next iteration. In the second step, the genome is scanned for pairwise marker interactions involving markers with significant additive effect from the previous step by considering likelihood ratio of a model with an interaction term to a model without such a term. We reapplied the first step of the analysis to the yeast dataset using the same folds we used for cross-validation (CV) for our ML methods. Additionally, we altered the CV procedure reported in the Nature paper (Bloom et al., 2013) as it was incorrect (the authors incorrectly identified QTLs on the training fold, but both fitted the model and obtained predictions on the test fold, which unfortunately overestimates the obtained *R*^2^ values, one of the pitfalls described by Wray et al. (2013)). We selected attributes and constructed the models only using the data in a training fold, with predictions obtained by applying the resulting model to the test fold.

The genomic BLUP model is a linear mixed-model similar to ridge regression but with a fixed, biologically meaningful penalty parameter. BLUP takes relatedness of the individuals in the study into account via genetic relatedness matrix computed from the genotypic matrix. The reason for choosing this method is that it (and its various extensions) is a very popular approach in genomic selection, and was the method applied in the original wheat paper (Poland et al., 2012). We used the R implementation in the RRBLUP package (Endelman, 2011).

### 3.3 Evaluation

The performance of all models was assessed using 10-fold and 5-fold cross-validation, for yeast and wheat, respectively and a train/test split for rice. Cross-validated predictions were collected across the folds and then used to calculate *R*^2^ (informally—proportion of variance explained by the model) in the usual manner (we call this measure cross-validated *R*^2^—cv*R*^2^). Wheat data set was small enough to repeat the CV procedure several times: we accumulated cv*R*^2^ across 10 runs on different fold selections and reported the average values. For rice, models were trained on 70% of the data and performance assessed on the remaining 30%.

## 4 Results

### 4.1 Overall Comparison of Methods

Tables 1, 2 and 3 summarise the cv*R*^2^ and *R*^2^ values for the standard statistical/machine learning methods, the Bloom GWAS method and BLUP for yeast, wheat and rice, respectively. The elastic net results are represented by the extremes, lasso, and ridge, as predictive accuracy appears to be a monotonic function of the elastic net penalty parameter *α* for all the datasets (see Figure 2).

**Table 2:** cv*R*^2^ for the five ML methods and for BLUP across 10 resampling runs. The best performance for each trait is in boldface.

| Trait/method | Lasso | Ridge | BLUP | GBM | RF | SVM |
| --- | --- | --- | --- | --- | --- | --- |
| Yield (drought) | 0.023 | 0.060 | 0.217 | 0.051 | 0.172 | **0.219** |
| Yield (irrigated) | 0.084 | 0.162 | 0.253 | 0.132 | 0.184 | **0.258** |
| TKW | 0.172 | 0.240 | 0.277 | 0.218 | 0.242 | **0.304** |
| DTH | 0.292 | 0.325 | 0.381 | 0.325 | 0.358 | **0.394** |

**Table 3:** *R*^2^ for the five ML methods and for BLUP using a train-test split for rice data. The best performance for each trait is in boldface.

| Trait/Method | Lasso | Ridge | BLUP | GBM | RF | SVM |
| --- | --- | --- | --- | --- | --- | --- |
| Culm Diameter | 0.142 | **0.191** | 0.190 | 0.145 | 0.163 | 0.182 |
| Culm Length | 0.529 | 0.567 | **0.568** | 0.559 | 0.539 | 0.516 |
| Culm Number | 0.205 | 0.232 | 0.233 | 0.188 | 0.222 | **0.247** |
| Grain Length | **0.387** | 0.375 | 0.381 | 0.371 | 0.361 | 0.383 |
| Grain Width | 0.474 | 0.508 | **0.511** | 0.466 | 0.435 | 0.500 |
| Grain Weight | 0.309 | 0.379 | 0.380 | 0.327 | 0.350 | **0.387** |
| Days to Heading | 0.677 | 0.693 | **0.698** | 0.669 | 0.657 | 0.636 |
| Ligule Length | 0.351 | 0.382 | 0.376 | 0.372 | 0.367 | **0.390** |
| Leaf Length | 0.335 | 0.406 | 0.405 | 0.407 | 0.388 | **0.414** |
| Leaf Width | 0.371 | 0.409 | 0.406 | 0.388 | 0.384 | **0.416** |
| Panicle Length | 0.352 | 0.399 | 0.399 | 0.383 | 0.388 | **0.416** |
| Seedling Height | 0.188 | 0.224 | **0.226** | 0.184 | 0.188 | 0.202 |

The yeast results show that there is at least one standard machine learning approach that outperforms Bloom and BLUP on all but 6 and 5 traits, respectively. In addition the mean advantage of the Bloom (BLUP) method on these 6 (5) traits is marginal: 1.8% (0.4%), with a maximum of 4.1% (1.5%), whilst the mean advantage of the standard ML techniques is 5.0% (5.5%), with a maximum of 14.2% (27.5%)—for SVM (GBM) on tunicamycin (maltose). (N.B. we did not re-run the second stage of the Bloom’s procedure, mining for pairwise marker interactions, but used the paper’s original results, so the actual cv*R*^2^ results for traits with interactions for Bloom’s method should be slightly lower than in Table 1). Across the ML methods the best performing method was GBM, which performed best in 26 problems. For 6 problems each lasso and the method of Bloom won. BLUP and SVM showed the best results for 5 and 3 traits, respectively. SVM on the whole across traits performed very similarly to BLUP but required time-consuming tuning.

The results for the wheat dataset paint quite a different picture: SVM performs the best for all traits (albeit with a marginal advantage for 3 out of 4 traits), followed closely by genomic BLUP. Both of the tree methods underperform compared to SVM and BLUP. The weakest method overall is lasso.

On the rice dataset, SVM also performs best for 6 of the 12 traits, genomic BLUP out-performs all other methods on 4 of the 12, and on a trait each, lasso and ridge perform best (see Table 3). As with the wheat dataset, the tree methods are also outperformed by BLUP and SVM, and the weakest method overall is also lasso. We hypothesise that BLUP’s ability to take genetic relatedness of the individuals into account gives it an advantage over the other two penalised regressions and also the two tree models.

The rest of this section is devoted to studying the performance of the five ML methods and BLUP on the yeast dataset in greater detail. As noted above, this form of dataset is arguably the cleanest and simplest possible.

### 4.2 Investigating the Importance of Mechanistic Complexity

The number of relevant attributes (markers, environmental factors), and the complexity of their interactions have an important impact on the ability to predict phenotype. We investigated how the mechanistic complexity of a phenotype impacted on the prediction results for the different prediction methods applied to the yeast dataset. Without a full mechanistic explanation for the cause of a phenotype, it is impossible to know the number of relevant attributes. However, in our yeast phenotype prediction data, which has no interacting environmental attributes, a reasonable proxy for the number of relevant attributes is the number of non-zero attributes selected by lasso regression (Figure 3). Therefore, to investigate the relationship between the number of markers chosen by the lasso and variance explained by the models we split the data into test (30%) and training (70%) sets, counted the number of non-zero parameters in the model fitted to the training set, and compared it to the model’s performance on the test set. We observed that the environments with the higher proportion of variance explained tend to have a higher number of associated non-zero attributes. Notable exceptions to this are the three (cadmium chloride, YPD-37C, and maltose) in the top-left part of the graph, which has an unusually high *R*^2^, but only a handful of associated non-zero attributes (only 6 markers for cadmium chloride). Notably, all three environments have a distinctive bimodal distribution that probably indicates that they are affected only by a few mutations.

**Fig. 3:**
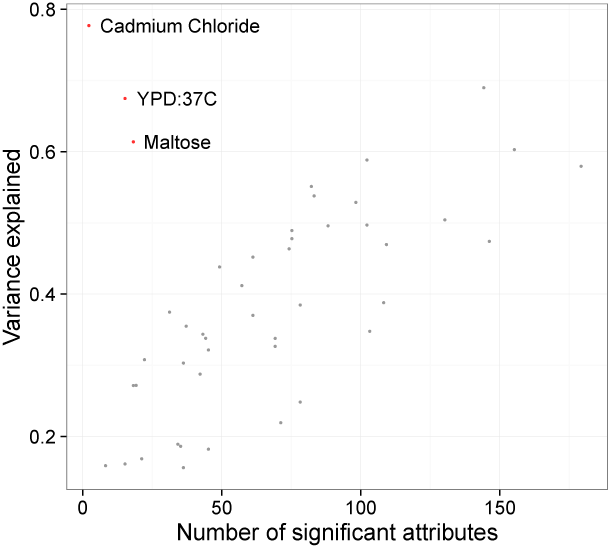
Non-zero attributes selected by lasso regression in training sample plotted against variance explained in the test sample (yeast).

We also wished to investigate how the complexity of the interactions of the attributes (in genetics the interaction of genes is termed ‘epistasis’) affected relative performances of the models. Figure 4 shows pairwise plots of relative performances of the seven approaches with red circles corresponding to those traits for which Bloom identified pairs of interacting attributes, and blue triangles corresponding to those for which no interacting attributes were found. We observed that lasso outperforms or matches (difference of less than 0.5%) Bloom’s method on those traits where no interactions were detected. This is true for all 22 such traits. Furthermore, we observed that the lasso on the whole also slightly outperforms GBM for some traits, and considerably outperforms random forest for the traits with no interactions. For traits with identified interactions boosted trees seem to show the best performance relative to results of Bloom and random forest; the former outperforms GBM for only 6 traits out of 46. Random forest seems to underperform compared to GBM and lasso, and it beats Bloom for only 14 traits with an average advantage of 1.9%.

**Fig. 4:**
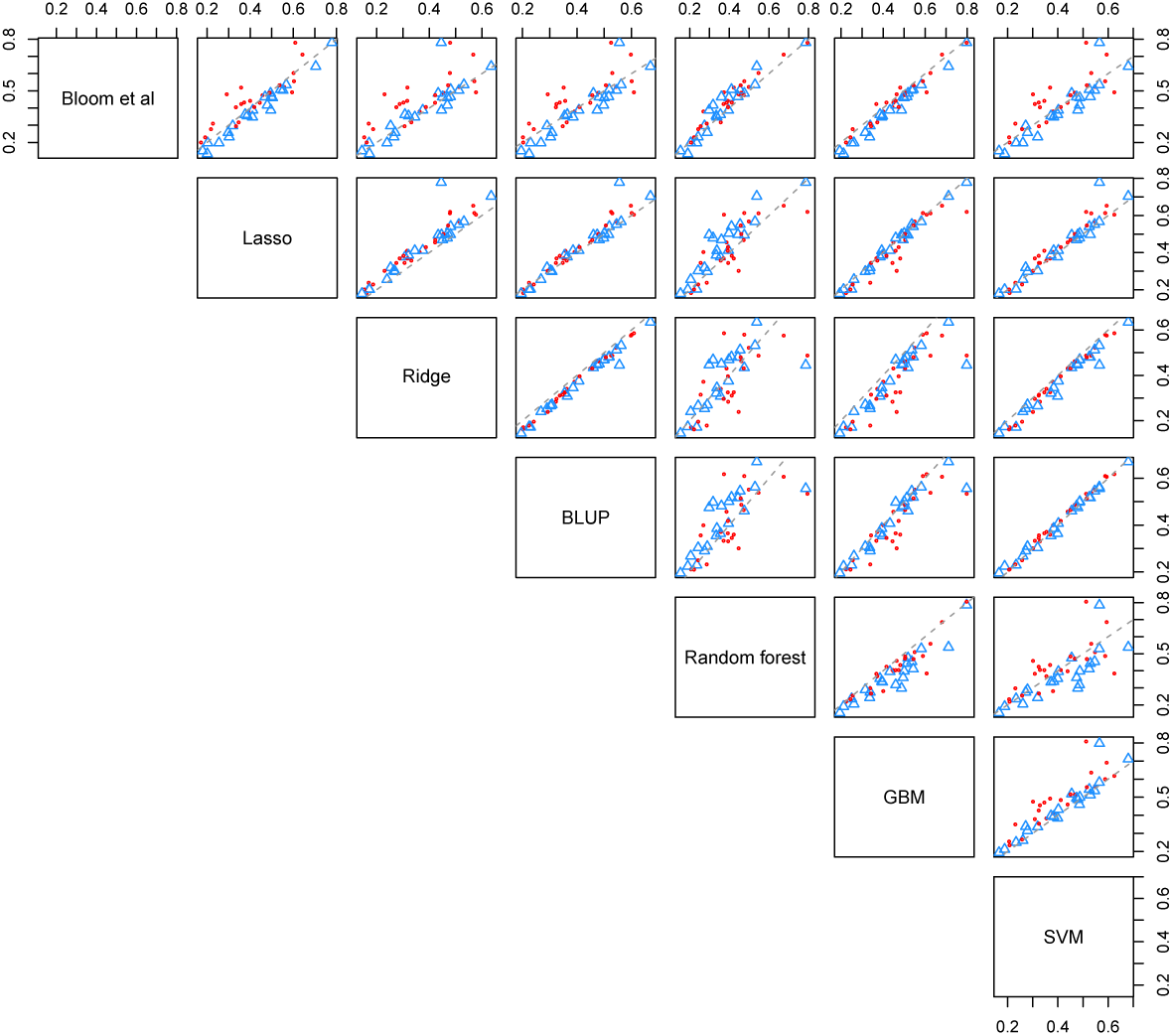
cv*R*^2^ for 10-fold cross-validation for different ML models, genomic BLUP and method of Bloom *et al.* applied to the yeast data. Red circles and blue triangles correspond to traits with interactions and no interactions (as identified by Bloom *et al.*), respectively.

Finally, we noted optimal tuning parameters chosen for GBM in each CV fold for each trait. In particular, we recorded tree interaction depth most frequently chosen across the 10 cross-validation folds (if two depths had equal frequency we took the smallest). Overall 1, 3 and 7 splits (corresponding to stumps, two-and three-way interaction trees, respectively) were chosen for 9, 18 and 19 traits, correspondingly. Looking closer, we noted that traits for which Bloom did not identify interacting attributes favoured stumps and shallow trees, while those with 2 or more interactions favoured deeper trees (7 splits)—in fact, 7 splits were identified as the optimal tree depth for all traits with more than two interactions. We conclude that optimal GBM tree depth might help draw conclusions about the structure and complexity of the underlying data.

### 4.3 Investigating the Importance of Noise in the Measured Phenotype

There is a great deal of noise in many real-world phenotype prediction problems. By noise, we mean both that the experimental conditions are not completely controlled, and the inherent stochastic nature of complex biological systems. Much of the experimental conditions noise is environmental (e.g., soil and weather differences for crop phenotypes, different lifestyles in medical phenotypes), and this cannot be investigated using the yeast dataset. However, in many phenotype prediction problems, there is also a significant amount of class noise. To investigate the importance of such class noise we randomly added or subtracted twice the standard deviation to a random subset of growth phenotype. We sequentially added noise to 5%, 10%, 20%, 30%, 40%, 50%, 75% and 90% of the phenotypic data for each trait and assessed performances of the statistical/machine learning methods on a test set (with training-testing split of 70%-30%). We repeated the procedure 10 times, each time selecting a different random subset of the data to add noise to. Figure 5a plots the ratio of variance explained using noisy phenotype versus original data versus proportion of noisy data added, averaged over the 10 runs. The results show a monotonic deterioration in accuracy with random forests performing the best, followed by GBM, BLUP, SVM, and lasso, with ridge regression trailing behind substantially. At about 20% of noisy data RF starts to outperform GBM in terms of average *R*^2^ (see Figure 5b).

**Fig. 5:**
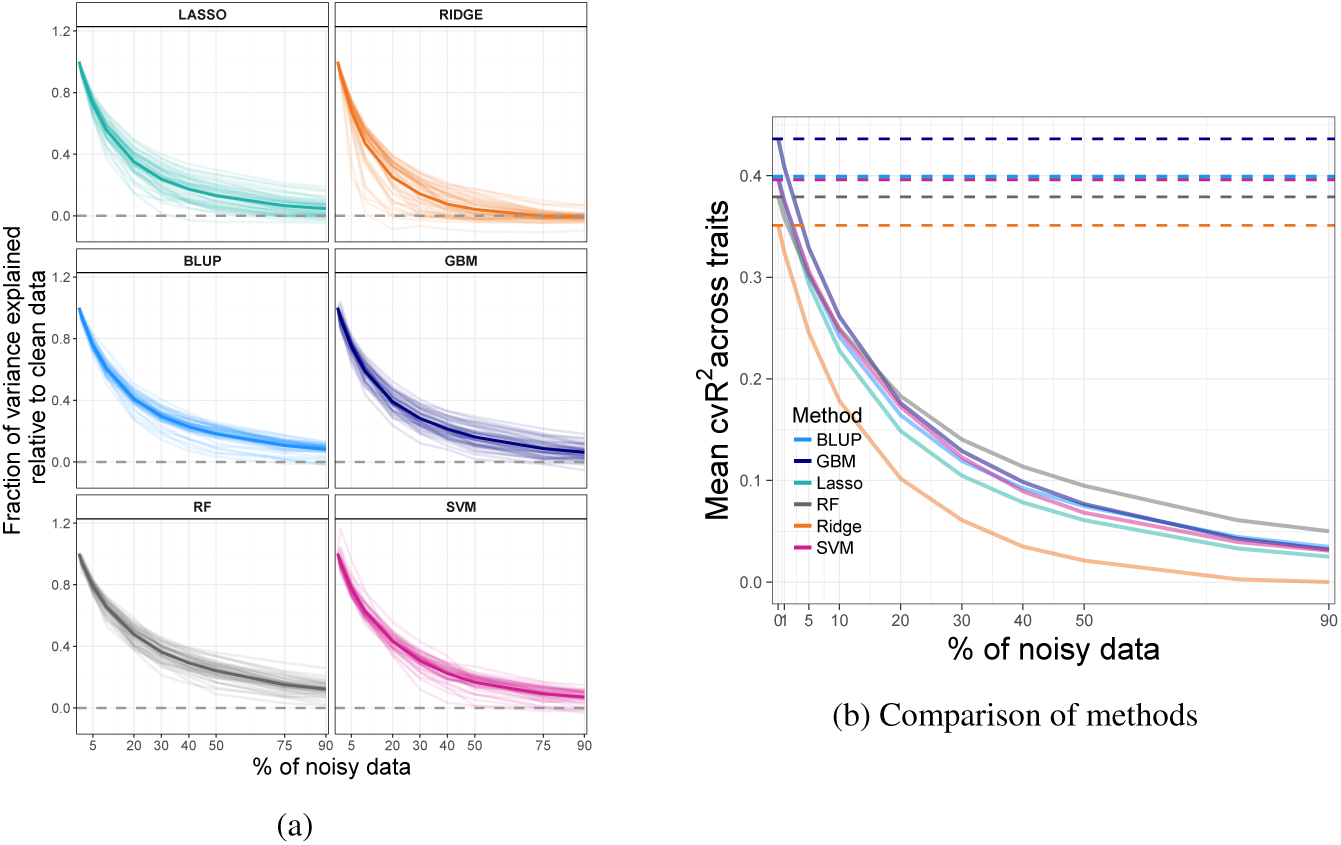
(a) Performance of ML and statistical methods under varying degrees of added class noise. The ratio of variance explained using noisy phenotype versus original data is plotted against proportion of noisy data added. The thick lines represent average value across all traits. (b) Comparison of methods on the absolute scale.

The algorithm underlying GBM is based on recursively explaining residuals of a model fit at the previous step, which might explain why it fares worse than RF under noise addition.

For all the methods and traits there seem to be very rapid deterioration in accuracy even for relatively small noise contamination (less than 10% of the data).

### 4.4 Investigating the Importance of Number of Genotypic Attributes

Another common form of noise in phenotype prediction studies is an insufficient number of markers to cover all sequence variations in the genome. This means that a genome is not fully represented, and other unobserved markers are present. To investigate this problem we sequentially deleted random 10%, 25%, 50%, 60%, 70%, 80%, 90%, 95% and 99% of the markers, and compared the performances of the five ML methods and BLUP on a test set (again with a training-testing split of 70%-30%). Figure 6a plots the ratio of variance explained using a reduced marker set versus variance explained using the full marker set versus proportion of markers deleted. Again these are average values over 10 runs with different random nested subsets of markers selected in each run. The statistical/machine learning methods, with the exception of ridge regression, lose minimal accuracy up until only 20% of the attributes remain, then undergo a rapid decline in accuracy after that. GBM and SVM seem to benefit from a reduced marker set for certain traits. BLUP’s performance is very consistent across the traits and seems to be affected by marker deletion the least. Absolute performance across all traits of different methods relative to each other remains unchanged for all levels of marker deletion (see Figure 6b).

**Fig. 6:**
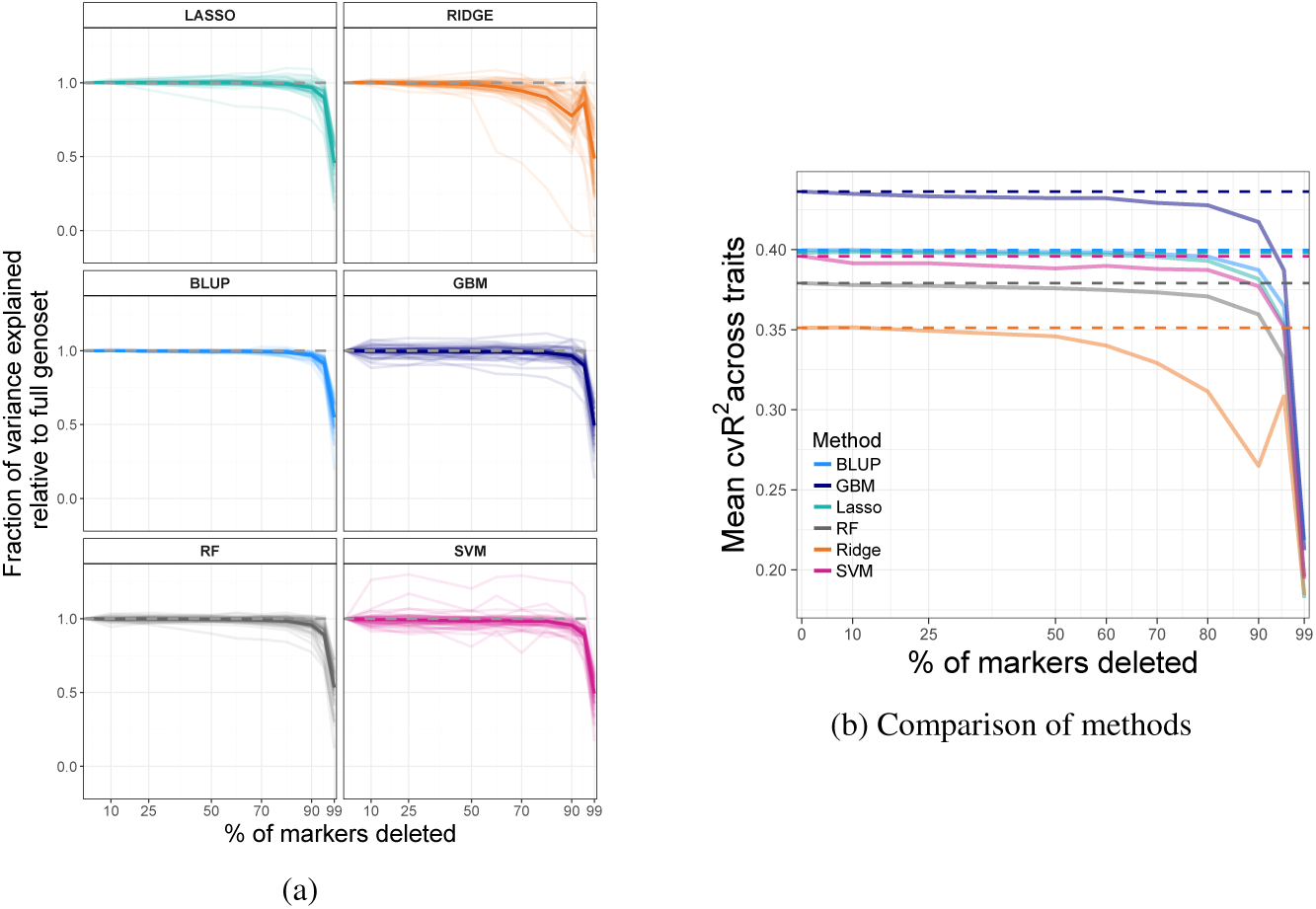
(a) Plots of the ratio of variance explained using a reduced marker set versus variance explained using the full marker set versus proportion of markers deleted for the five ML methods and BLUP. The thick lines represent average value across all traits. (b) Comparison of methods on the absolute scale.

The steepest dropping line in plots for lasso, ridge, GBM, and RF corresponds to cadmium chloride; this is most likely because only a handful of important markers are likely to be governing the trait (see Figure 3).

### 4.5 Investigating the Importance of Data Representation

For each yeast strain the marker data enabled us to recover the genomic structure formed by meiosis. We observed blocks of adjacent markers taking the same value, 1 or 0 (Figure 1a). As yeast has been fully sequenced and annotated (Cherry et al., 2012) we also know where the markers are relative to the open reading frames (ORFs), continuous stretches of the genome which can potentially code for a protein or a peptide,—’genes’. Knowledge of this structure can be used to reduce the number of attributes with minimal loss of information. We then investigated two ways of doing this. In the first, we generated a new attribute for each gene (in which there are one or more markers) and assigned it a value of 1 if the majority of markers sitting in it had a value 1, and 0 otherwise. In practice, we found that markers within each gene usually took on the same value for all but a handful of examples. Partially and fully overlapping genes were treated as separate. Markers within intergenic regions between adjacent genes were fused in a similar manner. Combining the gene and intergenic fused markers produced an alternative attribute set of 6,064 binary markers.

The second way we investigated the fusing of blocks of markers was to group markers in genes and their flanking regions. To do this, we divided the genome into regions, each of which contained one gene together with half of the two regions between it and the two neighbouring genes. Partially overlapping genes were treated separately, but genes contained entirely within another gene were ignored. Markers lying within gene regions formed in this manner were fused according to the dominant value within this gene. This produced an alternative set of 4,383 binary attributes.

We observed that the performance of the two alternative sets of genotypic attributes matched that of the full attribute set with the mean pairwise difference between any two attribute sets performance for each of the five ML methods, apart from ridge, not exceeding 0.5%. Ridge regression’s accuracy for some traits suffers considerably (e.g., cadmium chloride 5%, maltose 5-12%) when the reduced attribute sets were used. This indicates that most of the markers in blocks are in fact redundant as far as RF, GBM, SVM, and lasso are concerned.

### 4.6 Investigating the Importance of Number of Examples

In phenotype prediction studies there is often a shortage of data as it is expensive to collect. Traditionally obtaining the genotype data was most expensive, but increasingly data cost is being dominated by observation of the phenotype—with the cost of observing genotypes decreasing at super-exponential rate. To investigate the role of the number of examples we successively deleted 10%, 25%, 50%, 60%, 70%, 80% and 85% of all sample points and assessed performance of the six statistical/ML models (again on a test set with a train-test split of 70%-30% and 10 resampling runs). Figure 7a plots the ratio of variance explained using reduced data set versus original data versus proportion of data deleted. Figure 7b plots average performance over all traits for each method. Plots show that RF performed the most robustly. We note that by the time 50% of data is removed RF is outperforming lasso, BLUP, and SVM, and at the 80% mark it starts outperforming GBM (in absolute terms across traits, see Figure 7b). We notice that there is more variation across the traits in response to sample point deletion, as compared to addition of class noise and attribute deletion, where behaviour across the traits is more uniform (see Figures 5 & 6).

**Fig. 7:**
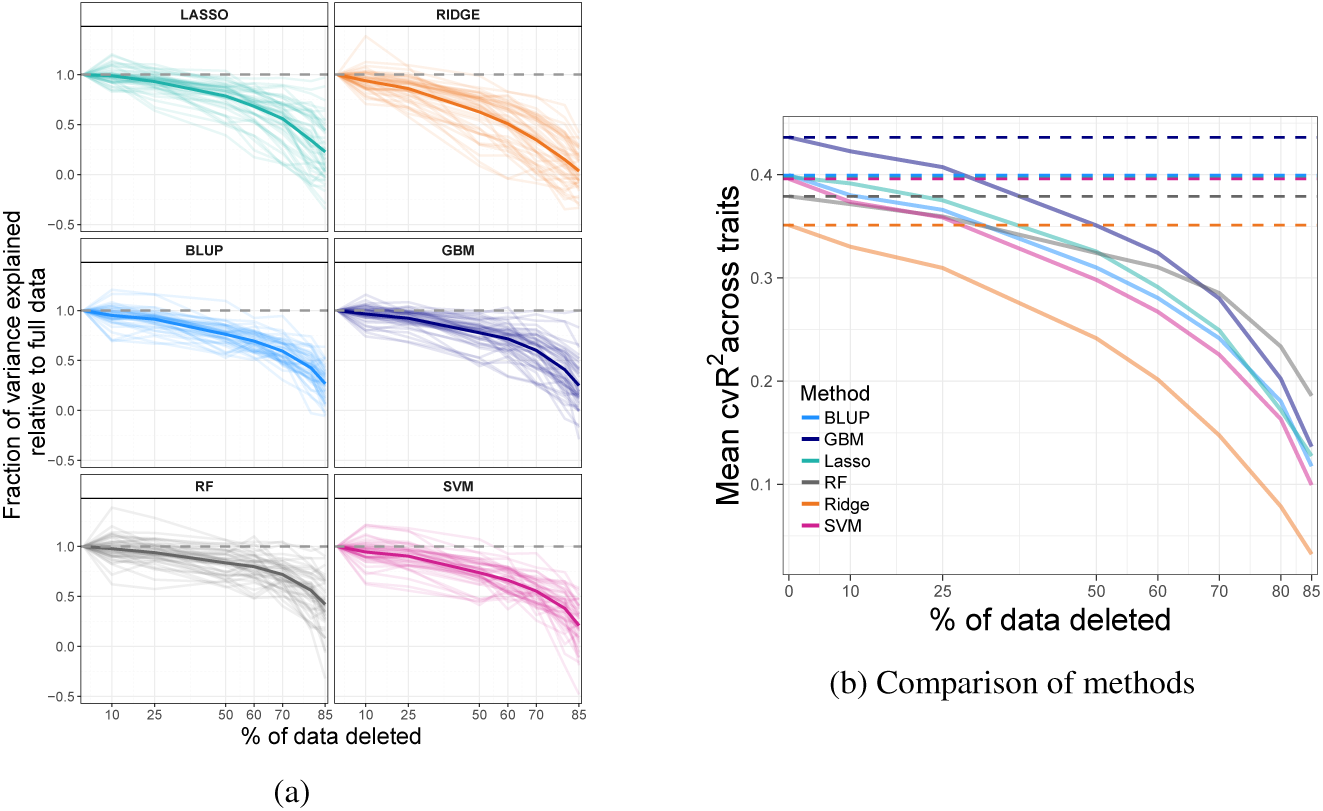
(a) Performance of ML methods and BLUP under sample points deletion. The ratio of variance explained using reduced dataset versus original is plotted against the proportion of data deleted. Thick lines represent average value across all traits. (b) Comparison of methods on the absolute scale.

### 4.7 Multi-Task Learning: Learning Across Traits

Rather than regarding the yeast dataset as 46 separate regression problems with 1,008 sample points in each, in the spirit of multi-task learning one might consider it as a single large regression problem with 46 ×1008 observations (in practice less due to missing values). One would then hope that a predictive model will learn to differentiate between different traits, giving accurate predictions regardless of the environments. Moreover, letting a model learn from several traits simultaneously might enhance predictive accuracy for individual traits through drawing additional information from other (possibly related) traits. We can think of this set-up as of a kind of transfer learning (Caruana, 1997; Evgeniou and Pontil, 2004; Ando and Tong, 2005). Most of the 46 traits are only weakly correlated (Pearson’s correlation) but there are several clusters of phenotypes with much higher pairwise correlations. Hence, on top of considering a regression problem unifying all 46 traits we also chose two smaller subsets of related traits with various levels of pairwise correlations:

a. Lactate, lactose, sorbitol and xylose: four sugar-related phenotypes with relatively high pairwise correlations (0.6-0.8).
b. Lactate, lactose, sorbitol, xylose, ethanol and raffinose: six sugar-related phenotypes with medium to high pairwise correlations (0.42-0.8).

Grouping these traits makes sense given yeast biology: xylose, sorbitol, lactose, and raffinose are all sugars, and plausible environments for yeast to grow in; ethanol is a product of yeast sugar fermentation; while lactate participates in yeast metabolic process. Hence it is not surprising that the six traits enjoy moderate to high pairwise correlations.

Combining several phenotype prediction problems into one results in each individual (yeast sample) having several entries in the input matrix and output vector—one for each trait. Additionally, we introduced an extra attribute, a categorical variable indicating which trait each sample corresponds to. We applied GBM and RF to this grouping approach. We show only random forest results, as GBM considerably underperformed compared to RF, and we did not apply SVM due to it being too computationally extensive for such a large problem. The performance was assessed by evaluation on a test set (30% of the data), which was an aggregation of test sets of individual traits used throughout the paper; this made comparing the new results to reference results obtained by training on separate traits easier.

We assessed the performance of RF on the two groups of traits above as well as on the group comprising all 46 traits. The three models were assessed for overall prediction accuracy as well as for prediction accuracy for each trait. We compared the latter values to reference fits, models trained and tested on each trait separately. Figure 8 below shows the results. One can see that 3 out of 4 traits that belong to both groups (*a*) and (*b*) (lactate, sorbitol, xylose) benefited greatly from grouped learning (blue and orange bars) whilst predictive accuracy for the two additional traits in group (*b*) (ethanol and raffinose) are substantially lower than reference. Moreover, adding these two traits reduced predictive accuracy for 3 out 4 of the original traits in group (*a*) (orange bars). Overall accuracy across all traits was 61% for group (*a*) and 52% for group (*b*).

**Fig. 8:**
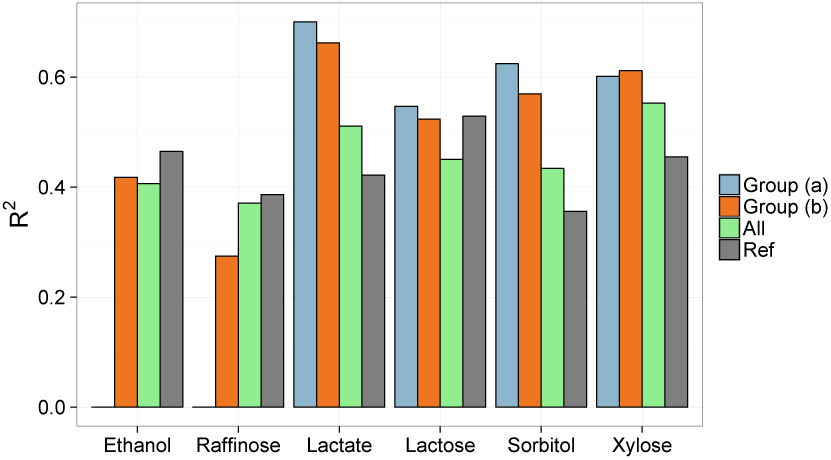
RF prediction accuracy (*R*^2^ on a test set) for individual traits when trained on group (*a*) (blue bars), group (*b*) (orange bars) and all of the 46 traits (green bars) compared to reference results (grey bars), when both learning and prediction was performed on individual traits.

For the full multi-task set-up using all 46 traits (green bars in Figure 8), the overall accuracy of all traits was just 20%. On average predictive accuracy for individual traits for this model was 22% lower than reference results (models trained and tested just on one trait). The accuracy for cadmium chloride, for example, dropped to just 0.4%. There were however 7 traits that benefited from grouped learning. Curiously these included lactate, sorbitol and xylose along with two other sugars trehalose and galactose (with an improvement of 5-10%). However, the accuracies for lactate, sorbitol and xylose were still lower than when these traits were considered as part of groups (*a*) and (*b*). It, therefore, seems that while combining multiple traits into a single regression problem indiscriminately might not, on the whole, improve overall or individual trait prediction accuracy, grouping carefully chosen traits with high pairwise correlation (perhaps advised by the organism’s biology) can be advantageous.

## 5 Discussion

We have demonstrated the utility of the application of machine learning methods on three phenotype prediction problems. The results obtained compared favourably to those obtained from state-of-the-art classical statistical genetics methods.

The yeast problem investigated has the simplest form of phenotype prediction problems. The data is also as clean and complete as is currently possible, and this enabled us to gradually degrade the data in various ways to better understand the factors involved in prediction success and to make it resemble other types of phenotype prediction problem. In the original clean yeast data, GBM performed best, with lasso regression and the method of Bloom *et al.* joint second best.

The wheat and rice problems are typical of crop genome selection problems. For this problem, SVM and BLUP were the best performing methods. We hypothesise that the success of BLUP is related to the population structure in this problem. Despite this success, BLUP does not optimally use population structure. Therefore, there is room to develop new machine learning methods that better use prior knowledge of population structure.

We investigated the role of the number of interactions between attributes. For yeast dataset traits with no interactions lasso proved to be the preferable method. We observed that GBM was the best method with traits with interacting attributes. In particular, traits with more than 2 interactions benefited deeper GBM trees. This suggests that complex interactions of order greater than two have utility in phenotype prediction problems, and that classical statistical genetics methods are mistaken to focus on univariate and bivariate interactions.

Out of the three types of noise we investigated class noise seem to be by far the most damaging to the prediction accuracy of all the methods. Of the machine learning methods, ridge regression’s performance deteriorated the most under various forms of noise, while random forest was the most robust method.

One important form of phenotype prediction problem that we have not studied is human disease associations studies. These problems typically differ from the problems investigated in several ways: there exists complex but unknown population structure, the environment of the examples is poorly controlled, and the phenotype investigated is a disease that may have a partly understood mechanism. These properties make such problems in some ways easier, and some ways harder, than the yeast and crop problems. We hypothesise that just as for the yeast, wheat, and rice datasets, the performance of off-the-shelf machine learning methods would compare favourably to those obtained from state-of-the-art classical statistical genetics methods.

There are a number of ways that machine learning methods could be developed for phenotype prediction problems. As mentioned in the introduction, the use of markers to describe genotypes is inefficient as it ignores their linear ordering, the type of sequence change, etc. One way to capture this information would be to use a relational representation (Getoor and Taskar, 2007). The marker representation also ignores all the prior biological knowledge that is available about the genes involved. For a gene, there may be known tens/hundreds/thousands of facts. This knowledge is highly relevant to determining whether a gene is mechanistically involved, but is often ignored by both ML and classical statistical genetics methods. To include this knowledge in prediction, a good approach would perhaps be to use a hybrid relational/propositional approach that uses relational data mining to identify useful attributes (King et al., 2001). A relational approach could also explicitly include prior background knowledge about the population structure of the examples.

For some GWAS problems the goal is to produce new organisms with desired properties, and an example of this is plant breeding where the goal is to produce plants with increased crop yield, resistance to drought, etc. This suggests an application of active learning. But this would require the development of new active learning methods that take into account the specific way (meiosis) that new examples (organisms) are produced.

To conclude, there is relatively little interaction between the machine learning and statistical genetics communities. This is unfortunate. Statistical genetics suffers from lack of access to new developments in machine learning, and machine learning suffers from a source of technically interesting, and societally important problems. We therefore hope that this paper will help bridge the differences between the communities and encourage machine learning research on phenotype prediction problems.

The original yeast data can be found at http://genomics-pubs.princeton.edu/YeastCrossBYxRM/, wheat data at https://dl.sciencesocieties.org/publications/tpg/abstracts/5/3/103, and rice data at http://snp-seek.irri.org/download.zul. Custom R code used to analyse the datasets can be found at: https://github.com/stas-g/grinberg-et-al-evaluation-of-ML-code.

This work was supported by BBSRC Grant BB/J006955/1.

